# ERBB-activated myofibroblastic cancer-associated fibroblasts promote local metastasis of pancreatic cancer

**DOI:** 10.1101/2022.12.05.519080

**Authors:** Gianluca Mucciolo, Joaquín Araos Henríquez, Sara Pinto Teles, Judhell S. Manansala, Muntadher Jihad, Wenlong Li, Eloise G. Lloyd, Priscilla S.W. Cheng, Giulia Biffi

## Abstract

Pancreatic ductal adenocarcinoma (PDAC) has a dismal prognosis. Cancer-associated fibroblasts (CAFs) are recognized potential therapeutic targets, but poor understanding of these heterogeneous cell populations has limited the development of effective treatment strategies. We previously identified TGF-β as a main driver of myofibroblastic CAFs (myCAFs). Here, we show that EGFR/ERBB2 signaling is induced by TGF-β in myCAFs through an autocrine process mediated by the ERBB ligand amphiregulin. Inhibition of this ERBB-signaling network in PDAC organoid-derived cultures and mouse models impacts distinct CAF subtypes, providing insights into mechanisms underpinning their heterogeneity. Remarkably, ERBB-activated myCAFs promote local PDAC metastasis in mice, unmasking functional significance in myCAF heterogeneity. Finally, analyses of other cancer datasets suggest these processes might operate in other malignancies. These data provide functional relevance to CAF heterogeneity and identify a potential target for preventing local tumor invasion in PDAC.

## INTRODUCTION

Pancreatic ductal adenocarcinoma (PDAC) is projected to be the second most common cause of cancer-related death by 2030^1^. PDAC is frequently lethal because it is often diagnosed late after patients have developed metastases. Dissecting metastatic mechanisms in PDAC and ways to prevent and treat this is therefore a priority. More than any other cancer, PDAC is characterized by an abundant, non-cancerous stroma that promotes cancer growth and treatment resistance. The majority of this stroma comprises a heterogeneous population of cancer-associated fibroblasts (CAFs)^2–10^, including molecularly and potentially functionally diverse myofibroblastic CAFs (myCAFs), inflammatory CAFs (iCAFs) and antigen-presenting CAFs (apCAFs)^6,9,11^. Understanding the pathways that maintain the identity and function of CAFs could unmask novel PDAC treatment approaches.

We previously identified interleukin 1 (IL-1) and transforming growth factor β (TGF-β) as the principal cancer-derived ligands that induce iCAF and myCAF formation, respectively^3^. While knowledge of pathways downstream of IL-1 signaling has revealed new iCAF treatment targets, pathways active in TGF-β-driven myCAFs are largely unknown.

## RESULTS

### TGF-β and PDAC organoid-conditioned media induce ERBB activation in myCAFs

TGF-β signaling is known to promote the formation and proliferation of PDAC myCAFs, but it is not known if this pathway serves other functions in these cells^3^. Therefore, we characterized receptor tyrosine kinases (RTK) phosphorylation following exposure of PDAC CAF precursor cells – pancreatic stellate cells (PSCs)^9,12^ – to TGF-β. Phosphorylated epidermal growth factor receptor (p-EGFR) and phosphorylated Erb-B2 receptor (p-ERBB2) were the most abundant RTKs activated upon TGF-β treatment, which was confirmed by western blotting in human PSCs (**Figures 1A-B and S1A**). Additionally, analysis of single-cell RNA-sequencing (scRNA-seq) datasets^6^ confirmed EGFR and ERBB2 expression in murine and human PDAC CAFs *in vivo* (**Figures S1B-C**).

**Figure 1.**
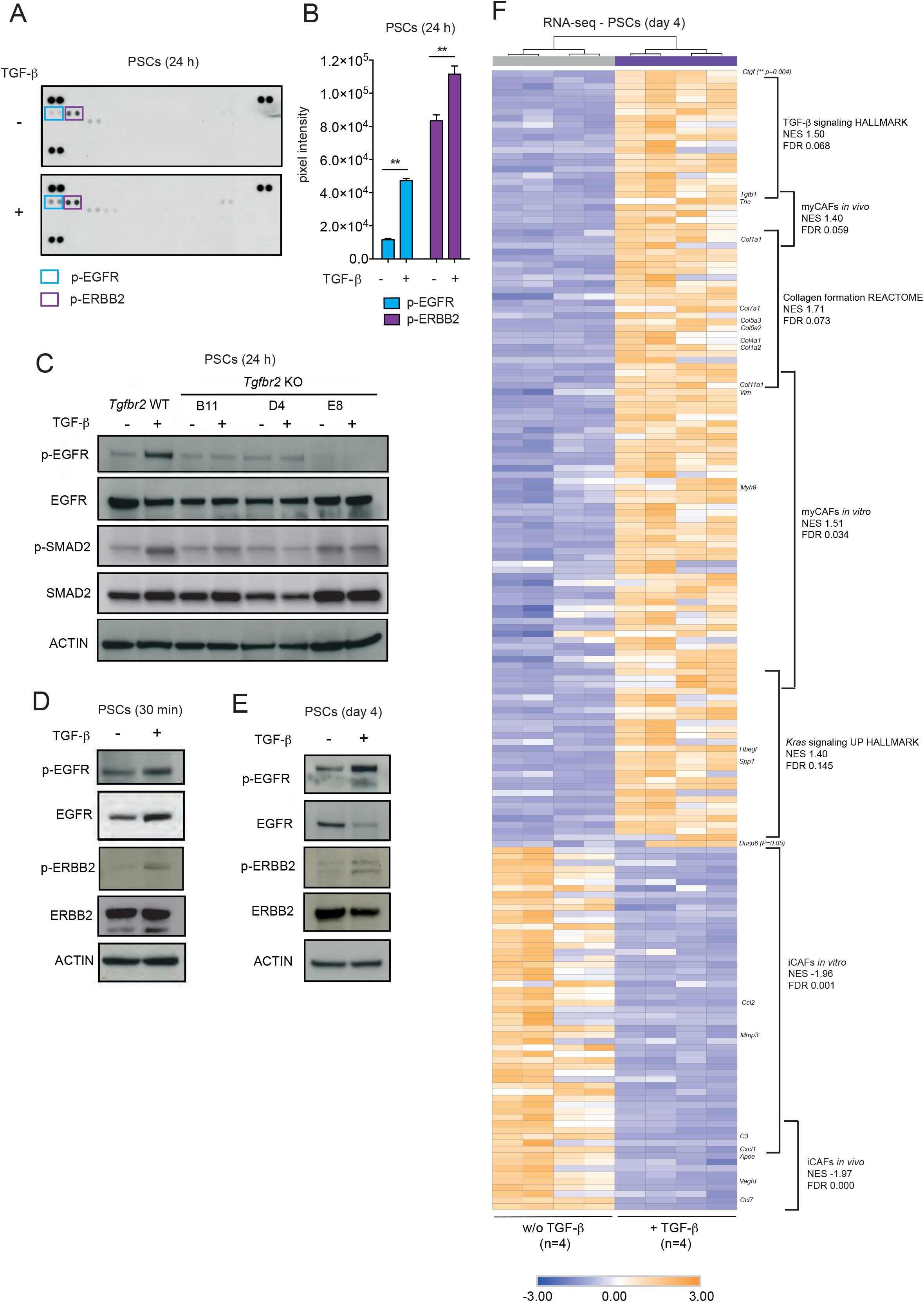
TGF-β induces ERBB activation in myCAFs. **(A)** Receptor tyrosine kinase (RTK) phosphorylation analysis of murine pancreatic stellate cells (PSCs) cultured for 24 h in Matrigel in control media (i.e. 5% FBS DMEM) with or without 20 ng/mL TGF-β. Blue and purple boxes highlight p-EGFR and p-ERBB2, respectively. **(B)** Quantification of p-EGFR and p-ERBB2 levels from A. Results show mean ± standard deviation (SD) of 2 technical replicates. **, *P* < 0.01, paired Student’s t-test. **(C)** Western blot analysis of p-EGFR, EGFR, p-SMAD2 and SMAD2 in murine *Tgfbr2* wild-type (WT) and knock out (KO) PSCs (3 clones from 3 different guide RNAs) cultured for 24 h in Matrigel in control media with or without 20 ng/mL TGF-β. ACTIN, loading control. **(D)** Western blot analysis of p-EGFR, EGFR, p-ERBB2 and ERBB2 in murine PSCs cultured for 30 min in Matrigel in control media with or without 20 ng/mL TGF-β. ACTIN, loading control. **(E)** Western blot analysis of p-EGFR, EGFR, p-ERBB2 and ERBB2 in murine PSCs cultured for 4 days in Matrigel in control media with or without 20 ng/mL TGF-β. ACTIN, loading control. **(F)** RNA-sequencing (RNA-seq) of PSCs cultured for 4 days in Matrigel in control media with or without 20 ng/mL TGF-β (n=4/group) showing selected genes and pathways enriched or downregulated upon TGF-β treatment. Color scheme of the heat map represents Z-score distribution. The myCAF and iCAF *in vitro* and *in vivo* signatures were obtained from Öhlund et al.^9^ and Elyada et al.^6^, respectively. Hierarchical clustering was determined by the top 50 most differentially expressed genes (DEGs). NES, normalized enrichment score; FDR, false discovery rate. See also **Figure S1**.

Deletion of TGF-β receptor II (TGFBR2) from PSCs blocked the induction of TGF-β responsive genes, TGF-β–dependent proliferation and activation of EGFR (**Figures 1C and S1D-F**). This suggests that TGF-β activates EGFR via its cognate receptor TGFBR2. Additionally, activation of EGFR and ERBB2 in PSCs was rapid, sustained and sensitive to TGF-β receptor I (TGFBR1) inhibition (**Figures 1D-E and S1G**). In keeping with its capacity to induce a myCAF cell fate, RNA-sequencing (RNA-seq) of TGF-β-treated PSCs revealed activation of a myCAF transcriptome and signatures associated with EGFR activation, including KRAS signaling and expression of *Dusp6*, a known target of the ERK pathway^13^ (**Figure 1F**).

TGF-β is expressed by PDAC cells *in vitro* and *in vivo* (**Figures S1H-J**). We showed previously that PDAC organoid-conditioned media (CM) activate SMAD2, a downstream member of the TGF-β pathway, in PSCs^3^. Therefore, we assessed whether PDAC organoid CM might activate EGFR/ERBB2 signaling in PSCs. In keeping with our studies of TGF-β, CM induced rapid and sustained activation of EGFR/ERBB2 in PSCs, which was blocked by the dual EGFR and ERBB2 receptor inhibitor (ERBBi) Neratinib (**Figure 2A**). Furthermore, PSCs treated with CM upregulated KRAS signaling, *Dusp6* expression and a PDAC myCAF-specific ERBB signaling signature, and these effects were blocked by Neratinib (**Figures 2B-D**). Targets in the myCAF-specific ERBB signature included regulators of cholesterol metabolism (**Figures 2C-D and Table 1**), and both myCAF-specific ERBB signature and cholesterol biosynthesis were also upregulated in TGF-β-activated myCAFs (**Figures S1K-L**). Finally, in keeping with a TGF-β /ERBB signaling network in myCAFs, Gene Set Variation Analysis (GSVA) of The Cancer Genome Atlas (TCGA) for PDAC (PAAD) identified a significant positive correlation between a patient-derived myCAF signature^6^, myCAF-associated TGF-β and Hedgehog (HH) signaling signatures, and EGFR activation (**Figure S1M**).

**Figure 2.**
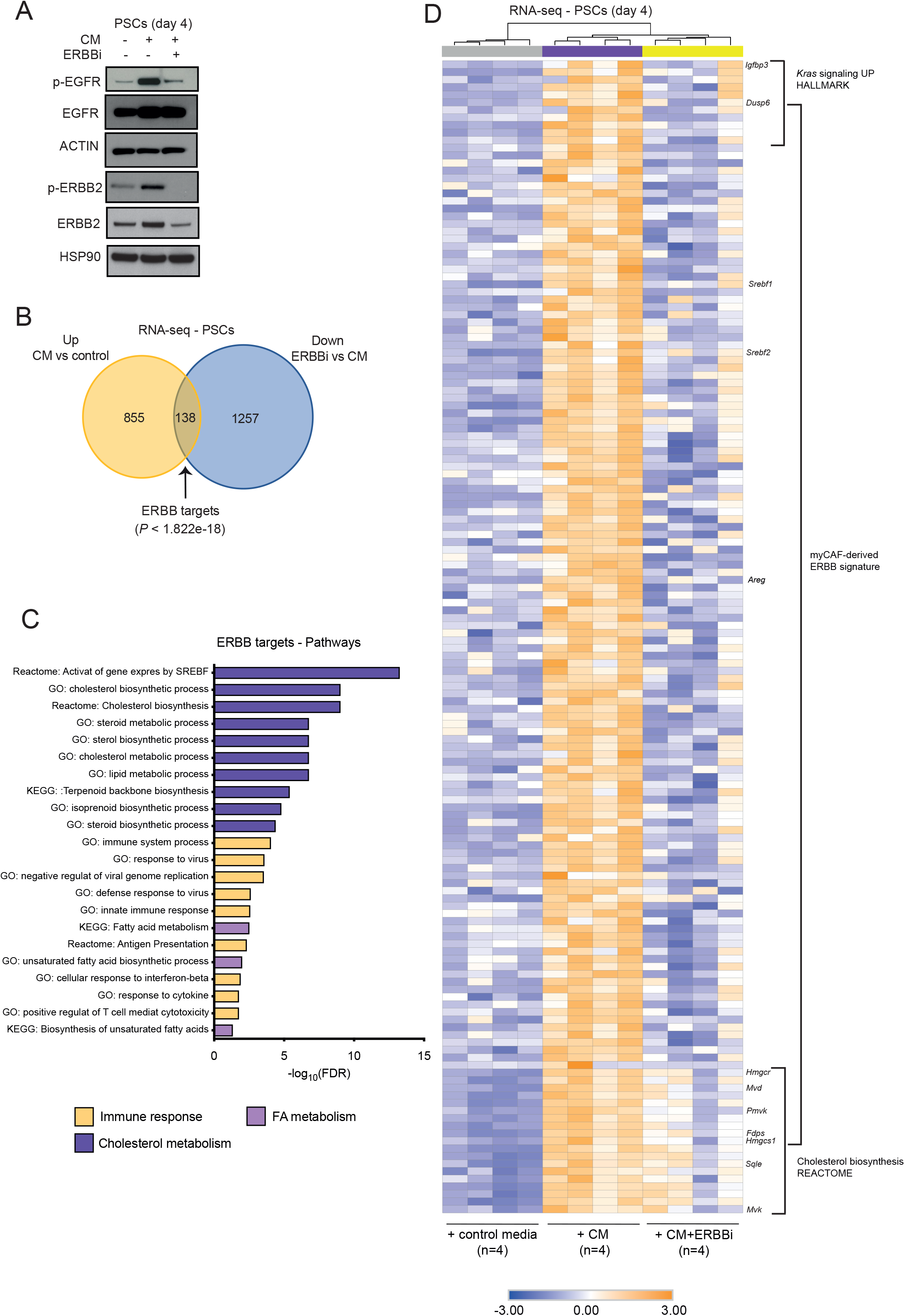
PDAC organoid-conditioned media induce ERBB activation in myCAFs. **(A)** Western blot analysis of p-EGFR, EGFR, p-ERBB2 and ERBB2 in murine PSCs cultured for 4 days in Matrigel in control media or PDAC organoid-conditioned media (CM) with or without 300 nM ERBB inhibitor (ERBBi) Neratinib. ACTIN and HSP90, loading controls. **(B)** Venn diagrams showing the overlap between significantly upregulated genes in PSCs cultured with PDAC organoid CM compared to PSCs cultured with control media and significantly downregulated genes in PSCs cultured with PDAC organoid CM + 300 nM Neratinib (ERBBi) compared to PSCs cultured with CM, as assessed by RNA-seq. The 138 genes present in both groups represent ERBB targets comprising the myCAF-derived ERBB signature. **(C)** Pathways found enriched in the myCAF-derived ERBB signature by DAVID analysis. Pathways were ranked by their significance (FDR<0.05) and significant terms (-log10 p value > 1.301) were highlighted. **(D)** RNA-seq of PSCs cultured for 4 days in Matrigel in control media or PDAC organoid-conditioned media with or without ERBBi (n=4-5/group) showing selected genes and pathways enriched or depleted. Color scheme of the heat map represents Z-score distribution. Hierarchical clustering was determined by the top 50 most DEGs. See also **Figure S1**.

Together, these data support a model in which PDAC cell-secreted TGF-β activates ERBB signaling in myCAFs in murine and human PDAC.

### A TGF-β-induced autocrine amphiregulin signaling mediates EGFR activation in myCAFs

Early activation of ERBB signaling in PSCs following treatment with TGF-β appeared to be mediated by increased receptor expression rather than ligand production (**Figures 1D and S2A**). To investigate how ERBB activation is sustained in myCAFs, we looked for expression of ERBB ligands in RNA-seq profiles of PSCs cultured with TGF-β or PDAC organoid CM. These RNA-seq profiles identified ERBB ligands, including amphiregulin (*Areg*) and heparin binding EGF-like growth factor (*Hbegf*), to be induced by TGF-β (**Figure 3A**). Real-time quantitative reverse transcription polymerase chain reaction (RT-qPCR) analysis confirmed TGF-β-induced expression of *Areg* and *Hbegf* in PSCs that was blocked by knockout of *Tgfbr2* or treatment with the TGFBR1 inhibitor (TGFBRi) A83-01 (**Figures 3B and S2B**). Moreover, neither EGFR deletion or inhibition - by Neratinib or the EGFR inhibitor Erlotinib (EGFRi) - completely ablated *Areg* and *Hbegf* expression by TGF-β (**Figures 3B and S2B-G**), validating *Areg* and *Hbegf* as candidate mediators of ERBB signaling in myCAFs. However, *Areg* was the only ERBB ligand significantly induced by both TGF-β and PDAC organoid CM (**Figure 3A**). Furthermore, only *TGFB1* and *AREG* expression were positively correlated in TCGA PAAD transcriptomes, suggesting AREG as the likely mediator of TGF-β-induced activation of ERBB signaling in myCAFs (**Figure S2H**). In addition, we confirmed upregulation of AREG protein by TGF-β in PSCs (**Figure 3C**).

**Figure 3.**
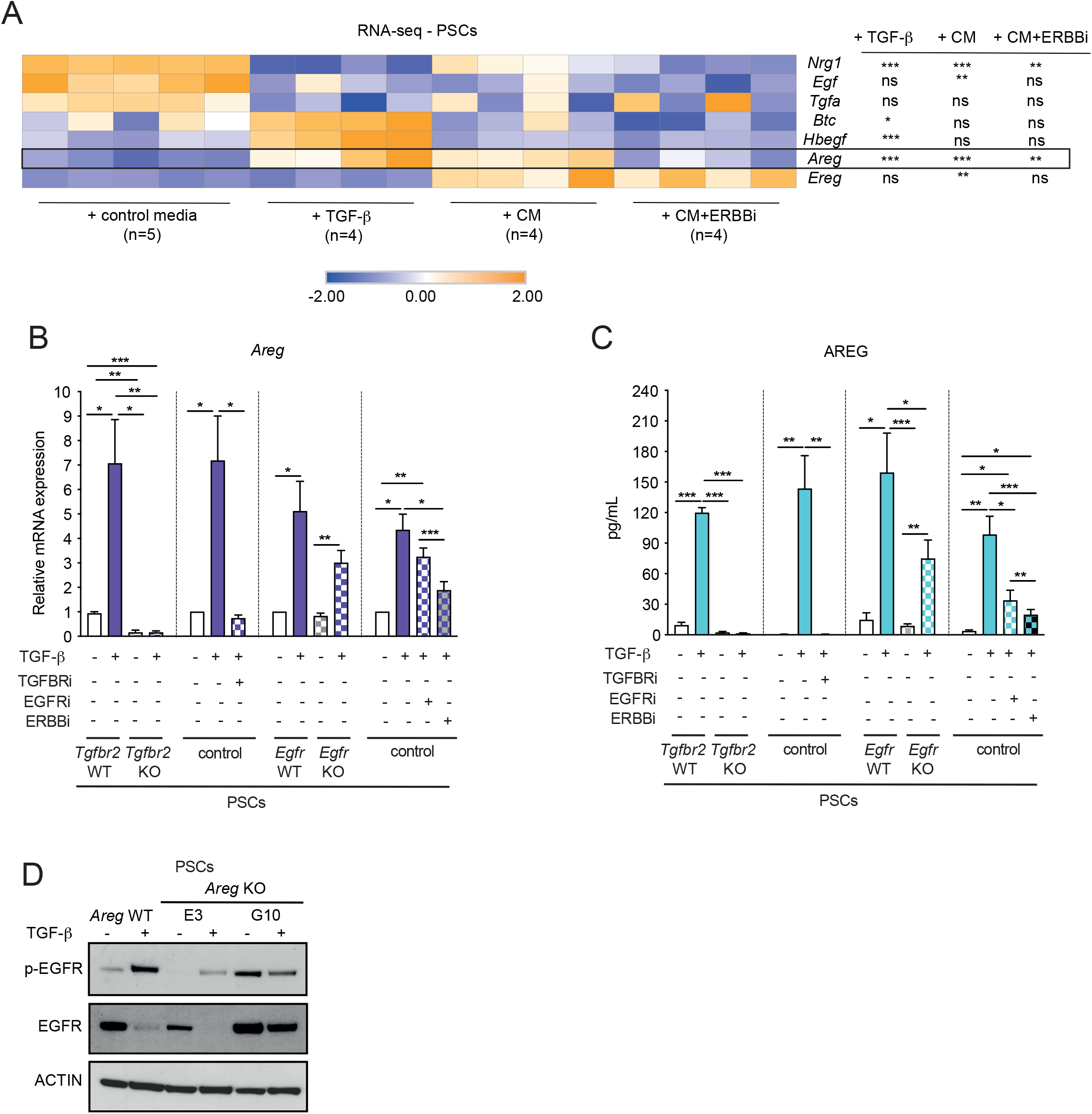
A TGF-β-induced autocrine amphiregulin signaling mediates EGFR activation in myCAFs. **(A)** RNA-seq expression of ERBB ligands (*Nrg1, Egf, Tgfa, Btc, Hbegf, Areg, Ereg*) in PSCs cultured for 4 days in Matrigel in control media, with 20 ng/mL TGF-β, PDAC organoid CM or CM with 300 nM Neratinib (ERBBi) (n=4-5/group). *, *P* < 0.05; **, *P* < 0.01, ***, *P* < 0.001, paired and unpaired Student’s t-test. Color scheme of the heat map represents Z-score distribution. **(B)** qPCR analysis of *Areg* in murine control (i.e. unmodified), WT (i.e. *Rosa26* KO), *Tgfbr2* KO or *Egfr* KO PSCs cultured for 4 days in Matrigel in control media with or without 20 ng/mL TGF-β in the presence or absence of 1 μM Erlotinib (EGFRi), 300 nM Neratinib (ERBBi) or 2 μM A83-01 (TGFBRi). Results show mean ± standard error of mean (SEM) of 4-8 biological replicates. *, *P* < 0.05; **, *P* < 0.01, ***, *P* < 0.001, paired and unpaired Student’s t-test. **(C)** ELISA of AREG from media of murine control, WT, *Tgfbr2* KO and *Egfr* KO PSCs cultured for 4 days in Matrigel in control media with or without 20 ng/mL TGF-β in the presence or absence of 1 μM Erlotinib (EGFRi), 300 nM Neratinib (ERBBi) or 2 μM A83-01 (TGFBRi). Results show mean ± SEM of 4-13 biological replicates. ***, *P* < 0.001, paired Student’s t-test. **(D)** Western blot analysis of p-EGFR and EGFR in murine *Areg* WT and KO PSCs cultured for 4 days in Matrigel in control media with or without 20 ng/mL TGF-β. ACTIN, loading control. See also **Figure S2**.

To test directly if AREG mediates the TGF-β-dependent activation of ERBB signaling in PDAC myCAFs, we deleted the gene from PSCs (**Figure S2I**). In agreement with our hypothesis, sustained EGFR activation induced by TGF-β was decreased in *Areg* KO PSCs relative to wildtype controls (**Figure 3D**). Notably, loss of AREG did not blunt the early activation of EGFR following TGF-β treatment, supporting the notion that this is a ligand-independent phenomenon (**Figure S2J**).

Thus, autocrine AREG mediates a sustained ERBB activation downstream of TGF-β signaling in PDAC myCAFs.

### Inhibition of ERBB signaling targets myCAFs *in vitro* and *in vivo*

To understand how ERBB activation impacts myCAFs, we first measured the proliferation of PSCs following TGF-β or PDAC organoid CM treatment in the presence of ERBB inhibitors. PSC proliferation was reduced significantly following both immediate or delayed (72 hours) exposure to the inhibitors without a detectable increase in apoptosis, suggesting that ERBB signaling mediates the TGF-β-dependent proliferation of PDAC myCAFs (**Figures 4A-B and S3A-F**). Accordingly, ERBB inhibition downregulated myCAF signatures (**Figure 4C**).

**Figure 4.**
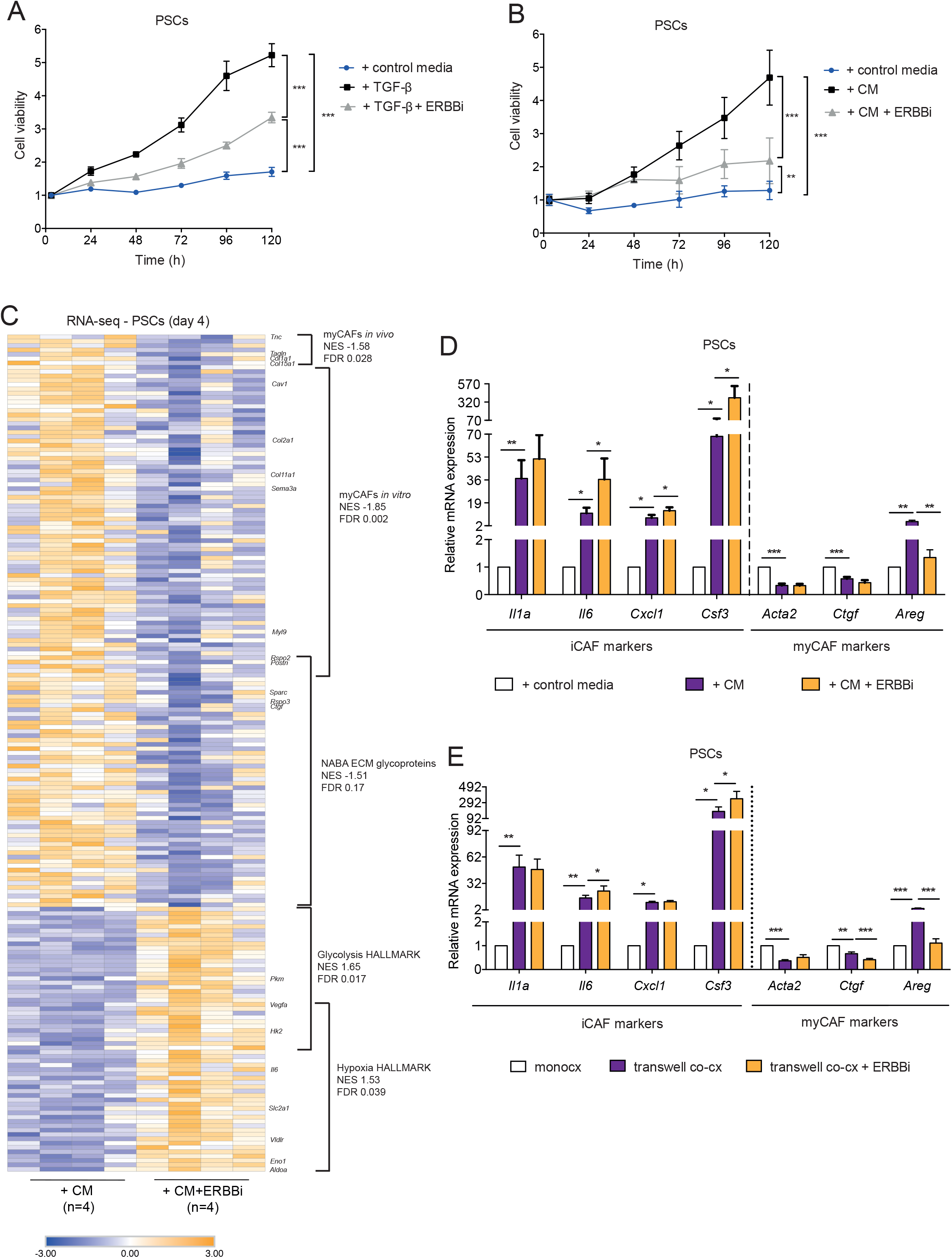
Inhibition of ERBB signaling impairs myCAF proliferation and signature in vitro. **(A)** Proliferation curves of murine PSCs cultured for 5 days in Matrigel in control media with or without 20 ng/mL TGF-β in the presence or absence of 300 nM Neratinib (ERBBi). Results show mean ± SD of 2 biological replicates (with 5 or 10 technical replicates respectively). ***, *P* < 0.001, unpaired Student’s t-test calculated for the last time point. **(B)** Proliferation curves of murine PSCs cultured for 5 days (120 h) in Matrigel in control media or PDAC organoid CM in the presence or absence of 300 nM Neratinib (ERBBi). Results show mean ± SD of 2 biological replicates (with 5 technical replicates each). **, *P* < 0.01; ***, *P* < 0.001, unpaired Student’s t-test calculated for the last time point. **(C)** RNA-seq analysis of PSCs cultured for 4 days in Matrigel in PDAC organoid CM (n=4) or CM in the presence of 300 nM Neratinib (ERBBi) (n=4) showing selected genes and pathways enriched or depleted upon ERBB inhibition. Color scheme of the heat map represents Z-score distribution. The *in vitro* and *in vivo* myCAF signatures were obtained from Öhlund et al.^9^ and Elyada et al.^6^, respectively. **(D)** qPCR analysis of *Areg*, and iCAF (*Il1a, Il6, Cxcl1, Csf3*) and myCAF (*Acta2, Ctgf*) markers in murine PSCs cultured for 4 days in Matrigel in control media, PDAC organoid CM or CM in the presence of 300 nM Neratinib (ERBBi). Results show mean ± SEM of 9 biological replicates. *, *P* < 0.05; **, *P* < 0.01; **\***\*\*, *P* < 0.001, paired Student’s t-test. **(E)** qPCR analysis of *Areg*, and iCAF (*Il1a, Il6, Cxcl1, Csf3*) and myCAF (*Acta2, Ctgf*) markers in murine PSCs cultured for 4 days in Matrigel in monoculture, in transwell culture with murine PDAC organoids or in transwell culture with murine PDAC organoids in the presence of 300 nM Neratinib (ERBBi). Results show mean ± SEM of 10 biological replicates. *, *P* < 0.05; **, *P* < 0.01; **\***\*\*, *P* < 0.001, paired Student’s t-test. See also **Figure S3**.

CAFs exist in different states and have been shown to at least partially interconvert upon pharmacological inhibition of pathways important for their formation^3–9^. Therefore, we looked to see if loss of the myCAF state might result in a reciprocal increase in iCAF fate. iCAF formation is dependent on JAK/STAT signaling and PDAC organoid CM treatment of PSCs induced iCAF-associated signatures that were blocked by JAK inhibition^3^ (**Figure S3G**). These iCAF-associated signatures, particularly those related to glycolysis and hypoxia, were increased upon ERBB inhibition in PSCs treated with PDAC organoid CM (**Figures 4C and S3H**), and qPCR analysis confirmed an upregulation of characteristic iCAF markers^3^ upon ERBB inhibition (**Figure 4D**). This same effect was observed when PSCs and PDAC organoids were cultured in transwell, even if PDAC organoid proliferation was affected by treatment with Neratinib (**Figures 4E and S3I**). Thus, ERBB inhibition preferentially targets PDAC myCAFs *in vitro*, leading to an enrichment in iCAFs.

To determine whether ERBB signaling inhibition affects CAF subtypes *in vivo*, we established orthotopic transplantation mouse models with PDAC organoids and treated tumor-bearing mice for 2 weeks with Neratinib (**Figures 5A and S4A**). Significant downregulation of p-EGFR levels and increased T cell abundance, which was previously reported following Erlotinib treatment^14^, confirmed effective targeting of the EGFR pathway (**Figures S4B-E**). In contrast to the impact of either TGFBR^3^ or HH^15^ inhibition on PDAC *in vivo*, Neratinib treatment did not reduce overall collagen deposition or the marker a-smooth muscle actin (aSMA), which are features of myCAFs (**Figures 5B-E**). Therefore, we looked to see if ERBB inhibition might differentially impact distinct myCAF subsets in tumors using our established flow cytometric quantification of Ly6C MHCII myCAFs, Ly6C^+^ MHCII iCAFs and LY6C MHCII^+^ apCAFs^6^ (**Figures S4F-G**). To further dissect heterogeneity among myCAFs, we also analyzed CAFs for CD90 (*Thy1*), which was previously shown to be highly expressed on a subset of myCAFs^15^ (**Figure S4H**). In agreement with our *in vitro* findings, myCAFs were significantly reduced upon Neratinib treatment, whereas iCAFs were significantly increased, altering the myCAF/non-myCAF ratio in tumors (**Figures 5F-G and S4I**). Notably, this effect was limited to CD90^-^ myCAFs since CD90^+^ myCAFs were unaffected by Neratinib treatment (**Figure 5H**).

**Figure 5.**
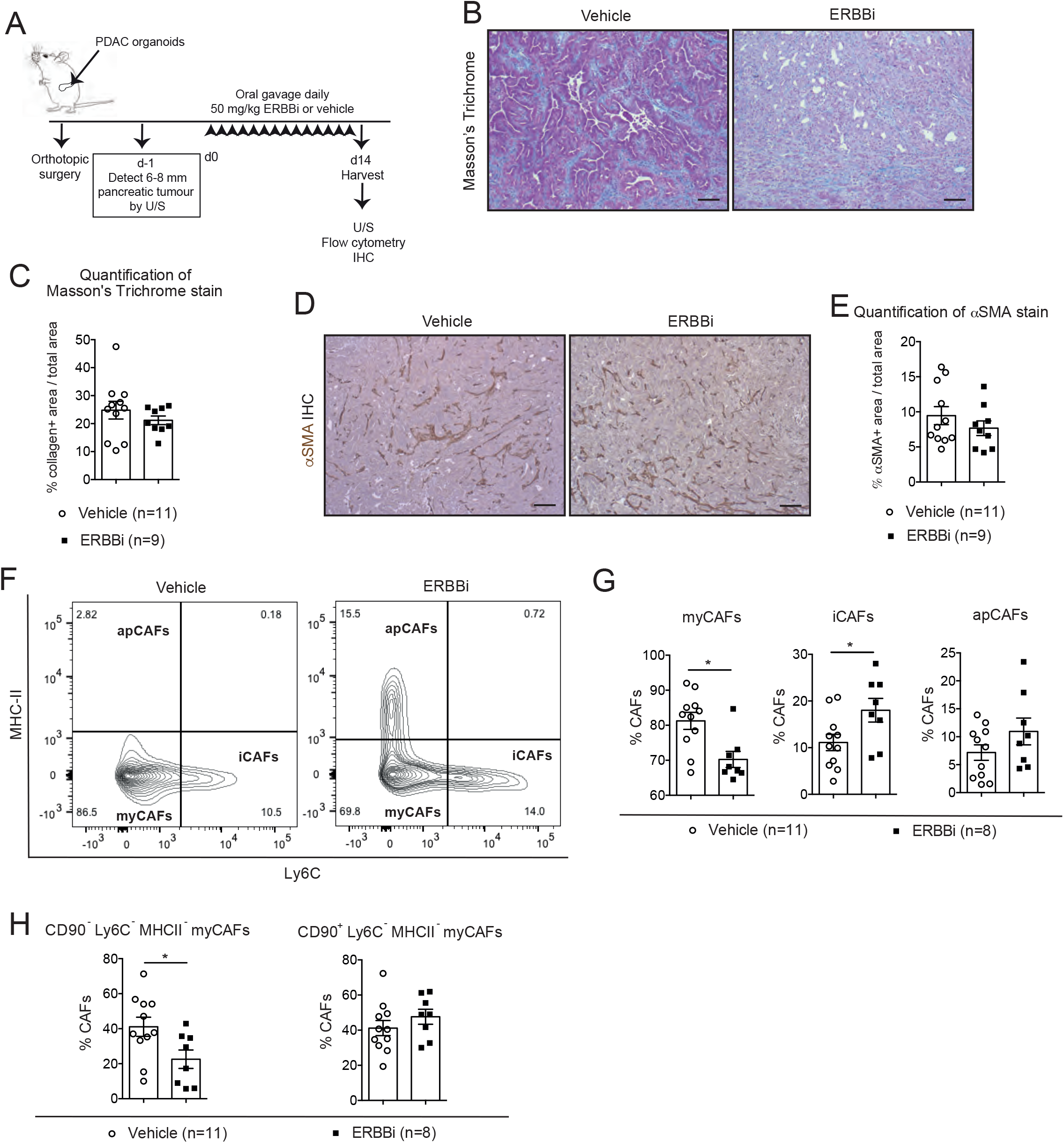
Inhibition of ERBB signaling targets myCAFs in vivo. **(A)** Schematic of 2-week treatment of tumor-bearing orthotopically grafted organoid-derived mouse models with 50 mg/kg ERBBi (Neratinib) or vehicle by daily oral gavage. U/S, ultrasound. **(B)** Representative Masson’s trichrome stain in 2-week vehicle- and ERBBi-treated tumors. Scale bars, 50 μm. **(C)** Quantification of Masson’s trichrome stain in 2-week vehicle-(n=11) and ERBBi-(n=9) treated tumors. Results show mean ± SEM. No statistical difference was found, as calculated by Mann-Whitney test. **(D)** Representative a-smooth muscle actin (aSMA) immunohistochemistry (IHC) in 2-week vehicle- and ERBBi-treated tumors. Scale bars, 50 μm. **(E)** Quantification of aSMA stain in 2-week vehicle-(n=11) and ERBBi-(n=9) treated tumors. Results show mean ± SEM. No statistical difference was found, as calculated by Mann-Whitney test. **(F)** Representative flow plots of myCAFs (Ly6C^-^ MHCII^-^), iCAFs (Ly6C^+^ MHCII^-^) and apCAFs (Ly6C^-^ MHCII^+^) from the PDPN^+^ parental gate in vehicle- and ERBBi-treated tumors. **(G)** Flow cytometric analyses of myCAFs (Ly6C^-^ MHCII^-^), iCAFs (Ly6C^+^ MHCII^-^) and apCAFs (Ly6C^-^ MHCII^+^) from the PDPN+ gate in vehicle-(n=11) and ERBBi-(n=8) treated tumors. Results show mean ± SEM. *, *P* < 0.05, Mann-Whitney test. **(H)** Flow cytometric analyses of CD90^-^ Ly6C^-^ MHCII^-^ myCAFs and CD90^+^ Ly6C^-^ MHCII^-^ myCAFs from the PDPN^+^ gate in vehicle-(n=11) and ERBBi-(n=8) treated tumors. Results show mean ± SEM. *, *P* < 0.05, Mann-Whitney test. See also **Figure S4**.

Together, these data provide insights into the heterogeneity of myCAFs and support the hypothesis that ERBB activation occurs in a subset of these CAF populations.

### ERBB-activated myCAFs promote local metastasis of PDAC

EGFR signaling in cancer cells has been previously described in PDAC tumorigenesis^16^. To investigate a potential role for EGFR-activated myCAFs in tumor progression, we established orthotopic transplantation mouse models of PDAC organoids alone or co-injected with *Egfr* WT or *Egfr* KO PSCs (**Figures 6A and S5A**). Detection by immunohistochemistry (IHC) of the cotransplanted PSCs, which are immortalized with the SV40 T antigen, confirmed the role of EGFR signaling in CAF proliferation, as observed *in vitro* (**Figures S5B-C and S2E-F**). While collagen deposition and aSMA levels were not altered across cohorts (**Figures S5D-G**), PDAC alone and PDAC+*Egfr* KO PSC tumors contained significantly fewer myCAFs and significantly more iCAFs compared to PDAC+*Egfr* WT PSC tumors, altering the myCAF/iCAF ratio (**Figures 6B and S5H-I**).

**Figure 6.**
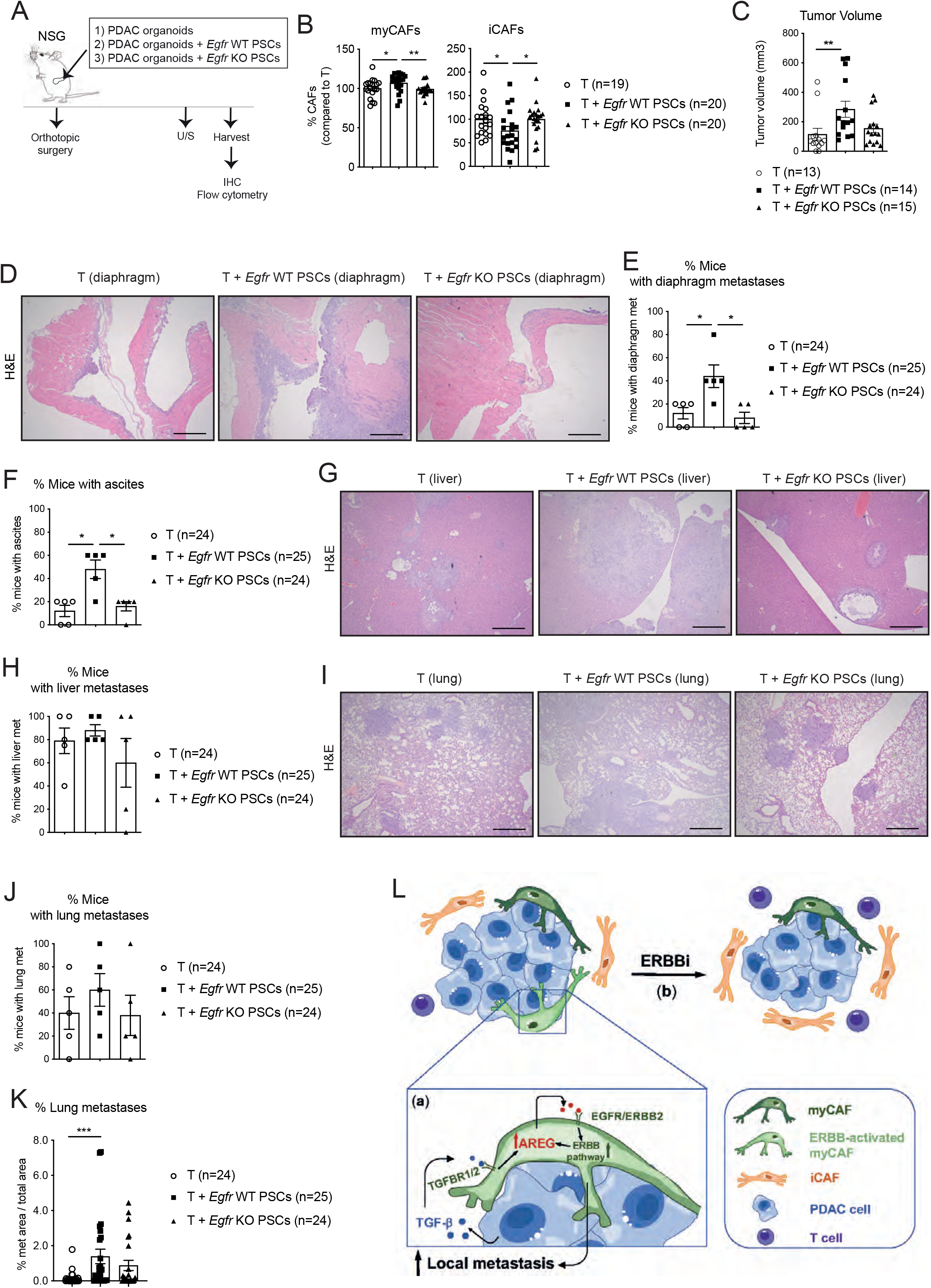
ERBB-activated myCAFs promote local metastasis of PDAC. **(A)** Schematic of experimental design of models in NSG mice derived by the transplantation of PDAC organoids (T) with or without *Egfr* WT or *Egfr* KO PSCs. U/S, ultrasound. **(B)** Flow cytometric analyses of myCAFs (Ly6C^-^ MHCII^-^) and iCAFs (Ly6C^+^ MHCII^-^) from the PDPN^+^ gate in tumors derived from the transplantation of PDAC organoids with or without *Egfr* WT or *Egfr* KO PSCs. Results show mean ± SEM (n=19-20/cohort). *, *P* < 0.05; **, *P* < 0.01, Mann-Whitney test. **(C)** Tumor volumes as measured by ultrasound of tumors derived from the transplantation of PDAC organoids with or without *Egfr* WT or *Egfr* KO PSCs. Results show mean ± SEM from 3 separate experiments (3-5 mice/cohort). **, *P* < 0.01, Mann-Whitney test. **(D)** Representative H&E stain of diaphragm tissues (with metastases) from mice transplanted with PDAC organoids with or without *Egfr* WT or *Egfr* KO PSCs. Scale bars, 200 μm. **(E)** Percentages of mice with diaphragm metastases in cohorts transplanted with PDAC organoids with or without *Egfr* WT or *Egfr* KO PSCs (n=24-25 mice/cohort). Results show mean ± SEM of 5 experiments (each with 4-5 mice per cohort per experiment). *, *P* < 0.05, Mann-Whitney test. **(F)** Percentages of mice with incidence of ascites in cohorts transplanted with PDAC organoids with or without *Egfr* WT or *Egfr* KO PSCs (n=24-25 mice/cohort). Results show mean ± SEM of 5 experiments (each with 4-5 mice per cohort per experiment). *, *P* < 0.05, Mann-Whitney test. **(G)** Representative hematoxylin and eosin (H&E) stain of liver tissues (with metastases) from mice transplanted with PDAC organoids with or without *Egfr* WT or *Egfr* KO PSCs. Scale bars, 200 μm. **(H)** Percentages of mice with liver metastases in cohorts transplanted with PDAC organoids with or without *Egfr* WT or *Egfr* KO PSCs (n=24-25 mice/cohort). Results show mean ± SEM of 5 experiments (each with 4-5 mice per cohort per experiment). No statistical difference was found, as calculated by Mann-Whitney test. **(I)** Representative H&E stain of lung tissues (with metastases) from mice transplanted with PDAC organoids with or without *Egfr* WT or *Egfr* KO PSCs. Scale bars, 200 μm. **(J)** Percentages of mice with lung metastases in cohorts transplanted with PDAC organoids with or without *Egfr* WT or *Egfr* KO PSCs (n=24-25 mice/cohort). Results show mean ± SEM of 5 experiments (each with 4-5 mice per cohort per experiment). No statistical difference was found, as calculated by Mann-Whitney test. **(K)** Quantification of metastatic burden (metastatic area/total area) in lung tissues of mice transplanted with PDAC organoids with or without *Egfr* WT or *Egfr* KO PSCs (n=24-25 mice/cohort). ***, *P* < 0.001, Mann-Whitney test. **(L)** Model illustrating how the ERBB pathway is activated in myCAFs downstream TGF-β signaling (a) and the effects of ERBB inhibition on PDAC CAF composition (b). See also **Figure S5**.

Remarkably, only tumors derived from PDAC+*Egfr* WT PSCs were significantly larger than those from PDAC alone (**Figure 6C**). Moreover, they generated significantly more diaphragm metastases and ascites than PDAC alone or PDAC+*Egfr* KO PSC tumors (**Figures 6D-F**). Additionally, mice with PDAC+*Egfr* WT PSC tumors had a significantly greater burden of lung metastases than those with PDAC tumors without PSCs, although the number of mice with evidence of liver or lung metastases were similar among cohorts (**Figures 6G-K and S5J**).

Altogether, these data identify a previously unappreciated functional complexity of myCAFs, showing that ERBB-activated myCAFs promote local metastasis of PDAC (**Figure 6L**).

### EGFR activation occurs in myofibroblastic CAFs in various malignancies

As PDAC CAFs share features with CAF subtypes in other malignancies^4^, we investigated the broader impact of our findings among malignancies in which ERBB inhibition is an established therapeutic strategy^17^. Similar to what observed in the PDAC dataset (**Figure S1M**), GSVA analysis of TCGA breast cancer BRCA dataset using a myofibroblastic CAF signature^18^ identified pathways known to be activated in myCAFs, such as TGF-β and HH signaling, and confirmed a positive correlation between the myofibroblastic CAF signature and EGFR activation (**Figure S6A**). Additionally, similar to what we found in TCGA PAAD (**Figure S2H**), analysis of TCGA BRCA and lung cancer LUAD datasets showed a positive correlation between *TGFB1* expression and expression of *AREG*, as well as of other myCAF markers (**Figures S6B-C**). Finally, TGF-β treatment induced *Areg* and *Dusp6* expression and EGFR activation in mouse pulmonary fibroblasts (**Figures S6D-F**).

Together, these analyses suggest that EGFR activation occurs also in TGF-β-dependent myofibroblastic CAFs of other malignancies and could be directly affected by ERBB-targeting strategies, as shown in PDAC.

## DISCUSSION

We reveal a previously unknown role for EGFR activation in a population of PDAC CAFs. Our data show that TGF-β induces AREG expression by PDAC myCAFs, triggering an autocrine EGFR/ERBB2 response. Since ERBB blockade downregulates AREG expression, this suggests a positive feedback loop within this ligand/receptor network. This network appears to fine tune the balance of CAF cell fates, favoring a myCAF relative to iCAF phenotype. This effect appears to be restricted to a subpopulation of ERBB-activated myCAFs that promotes local PDAC metastasis in mice (**Figure 6L**). We thereby unmask a new mechanism by which the cancer-CAF cross-talk regulates PDAC myCAF heterogeneity and metastasis.

Phospho-EGFR has been previously detected in non-cancer cells in a *Kras^G12D^*; *Egfr^KO^* mouse model of PDAC^16^, and AREG has been previously shown to promote sustained EGFR activation in homeostasis and inflammation^19–21^. Our work supports a role for CAF-autocrine AREG signaling in sustaining EGFR activation in TGF-β-driven myCAFs. However, AREG is also secreted by cancer and/or immune cells. For example, it has been demonstrated that regulatory T cell (Treg) depletion leads to loss of myCAFs in PDAC^22^. Although this is likely dependent on TGF-β, Tregs also produce AREG^23^, whose reduction upon Treg depletion may also be involved in the observed reduction in myCAFs. Finally, since both EGFR and ERBB2 are activated upon culture with TGF-β, and Neratinib treatment led to a more pronounced downregulation of *Areg* compared to Erlotinib, we speculate that the EGFR/ERBB2 heterodimer is involved in AREG induction and ERBB signaling in PDAC myCAFs.

While we identified a tumor-promoting role of ERBB-activated myCAFs, previous work proposed a tumor-restraining role of myCAFs^24^’^25^, largely attributing this to myCAF-mediated collagen deposition. We show that ERBB inhibition does not impact collagen abundance, likely because the ERBB signaling pathway appears active only in a subset of myCAFs. Together, these observations highlight the complexity of CAF populations and the need to further understand their molecular and functional heterogeneity.

CAFs appear to promote metastases through various mechanisms. They can drive cancer cell aggressiveness by secreting ligands^26–29^, increase their viability and provide early growth advantage at secondary sites by co-migrating with them^30,31^, or exert force to drive cancer cell collective migration and invasiveness^32–34^. Future work will be required to fully dissect the mechanism behind the promotion of local PDAC metastasis by ERBB-activated myCAFs.

In agreement with published data^14^, we observed an increase in T cell abundance following ERBB inhibition. CAF populations have been implicated in regulating the immune microenvironment^4^, and we show that an ERBB signaling network contributes to CAF heterogeneity. Together, these data underscore the complex interplay between distinct CAF subtypes, immune cells and PDAC progression. Extensive future work will be required to fully understand how these processes operate to regulate the PDAC microenvironment and PDAC metastasis.

As recent studies suggest that EGFR inhibition in PDAC may be helpful in combination with immunotherapies^14^, benefit *EGFR* WT cases^35^ and revert resistance to KRAS^G12C^ inhibitors^36^, our study could be clinically relevant for PDAC patients. Additionally, our observations could have a broader impact, as our analyses suggest that activation of ERBB signaling also occurs in myofibroblasts of breast cancer and lung cancer, in which the ERBB/EGFR pathway is more commonly inhibited in the clinic. Similarly, previous work has implicated EGFR activation and AREG upregulation in myofibroblasts in liver and pulmonary fibrosis^37–40^. Therefore, AREG/ERBB signaling may be common to numerous fibrotic diseases in which myofibroblasts play major roles.

Our study reveals ERBB signaling as a TGF-β-dependent pathway active in PDAC myCAFs; highlights a previously unappreciated effect of ERBB signaling inhibition on the PDAC stroma that might also operate in other malignancies; and identifies a role for ERBB-activated PDAC myCAFs in promoting local metastasis.

## ACKNOWLEDGEMENTS

The authors would like to thank the BRU, Genomics, Bioinformatics, Flow cytometry, Pregenome editing and Histology core facilities at the Cancer Research UK Cambridge Institute (CRUK-CI). This work was mainly supported by a Cancer Research UK institutional grant (A27463), which also supported G.B., S.P.T and J.S.M. G.B. is recipient of a UKRI Future Leaders Fellowship, which also supports W.L., a Pancreatic Cancer Research Foundation grant and a US Department of Defense PCARP grant, which support G.M. and J.S.M., a NCI-CRUK Cancer Grand Challenge grant, which supports M.J., and a Pancreatic Cancer UK Future Leaders Academy grant, which supports P.S.W.C. J.A.H. is supported by a Harding Distinguished Postgraduate Programme PhD studentship (Cambridge Trust). E.G.L. is supported by a MRC Doctoral Training Grant. The results shown here are in part based on data generated by the TCGA Research Network (http://www.cancer.gov/tcga).

## AUTHOR CONTRIBUTIONS

G.M. and J.A.H. designed the experiments and conducted the experiments. S.P.T, J.S.M., M.J., W.L., E.G.L. and P.S.W.C. conducted the experiments. G.B. designed the experiments, conducted the experiments and wrote the paper.

## TABLE LEGENDS

**Table 1. Differential expression analysis of PSCs by RNA-seq.** Significant protein coding differentially expressed genes (DEGs) (FDR < 0.05) between TGF-β treated PSCs and PSCs in 5% FBS DMEM (DEGs TGF-β vs Control) or CM-treated PSCs and PSCs in 5% FBS DMEM (DEGs CM vs Control) are summarised in this table. Additionally, significant protein coding DEGs of CM-treated PSCs in the presence of Neratinib in contrast to CM-treated PSCs in the absence of Neratinib (DEGs ERBBi vs CM) are shown. Moreover, we include a list of 138 ERBB target genes (ERBB signature) common between upregulated DEGs CM vs Control (logFC > 0) and downregulated DEGs ERBBi vs CM (logFC < 0).

## METHODS

### Mouse models

Males and females C57BL/6J (strain number 632) and NSG mice (strain number 614) were purchased from the Charles River Laboratory. All animals are housed in accordance with the guidelines of the UK Home Office “Code of Practice for the Housing and Care of Experimental Animals”. They are kept behind strict barriered housing, which has maintained animals at a well-defined microbiological health status. This accommodation precludes access by wildlife, including rodent and insect vectors, and is free of infestation with ectoparasites. All animals are health screened every 3 months according to the FELASA guidelines (FELASA 2002). All animals are fed expanded rodent diet (Labdiet) and filtered water ad libitum. Environmental enrichment includes nesting material, structures for three-dimensional use of the cage and an area to retreat, and provision of chew blocks. All animal procedures and studies were reviewed by the CRUK-CI AWERB, approved by the Home Office and conducted under PPL number PP4778090 in accordance with relevant institutional and national guidelines and regulations.

### Orthotopic transplantation models

Orthotopic injections were conducted as previously described^3^. Briefly, single cells (10,000 cells/mouse) prepared from organoid cultures (female T69A or male T6-LOH) were resuspended as a 40 μL suspension of 50% Matrigel in PBS and injected into the pancreas of 8-10-week-old mice with or without 10,000 (1:1) *Egfr* WT or KO PSCs. Pancreatic tumors in NSG mice were only imaged once using the Vevo 2100 Ultrasound at two different orientations with respect to the transducer. Tumor volumes were measured at two angles using the Vevo LAB software program (version 5.7.0).

### Neratinib treatment

Pancreatic tumors in C57BL/6J mice were imaged prior to enrolment (day-1) and at endpoint (day 14) using the Vevo 2100 Ultrasound at two different orientations with respect to the transducer. Mice with tumor diameters of 6 to 8 mm were randomized and enrolled 1 day after scanning. Tumor volumes were measured as above, and growth rate was measured by dividing the volume at day 14 for the volume at day-1. The ERBB inhibitor Neratinib was prepared daily as a suspension in 0.1% Tween80, 0.5% hydroxyl propyl methyl cellulose in sterile water. Mice were administered vehicle or 50 mg/kg of Neratinib for 14 days, once a day via oral gavage.

### Cell lines and cell culture

Mouse PSCs (SV40-immortalised) and tumor pancreatic organoid lines were previously described^9,41^. Human PSCs (SV40-immortalised) were purchased from ScienCell (3830). Mouse PSCs and human PSCs were cultured in DMEM (41966029; Gibco) containing 5% FBS. Mouse pulmonary fibroblasts from C57BL/6 were purchased from Caltag Medsystems (SC-M3300-57), SV40-immortalised and cultured in fibroblast medium basal (SC-2301-B, Caltag Medsystems) with 10% FBS. All cells were cultured for no more than 30 passages at 37C with 5% CO2. For conditioned media experiments, tumor organoids were cultured for 3 to 4 days in DMEM with 5% FBS (i.e. control media). For transwell cultures, organoids were plated on top of transwell membranes (82051-572; VWR) with PSCs growing in Matrigel (356231 and 356230; Corning) in 24-well plates in DMEM with 5% FBS. Cell line authentication was performed at the CRUK-CI for the murine PSCs. Mycoplasma testing is performed weekly, and each cell line is tested prior to freezing.

### *In vitro* cell treatments

PSCs were treated in Matrigel in 5%FBS DMEM with 20 ng/mL human TGFβ1 (T7039; Sigma), 300 nM Neratinib (S2150; Selleckchem), 1 μM Erlotinib HCl (S1023-SEL; Stratech Scientific Ltd), 2 μM A83-01 (2939; Tocris Bioscience), 10 ng/mL murine EGF (PMG8043; Thermofisher Scientific) for as long as specified in the figure legends.

### *Tgfbr2, Egfr, Areg* CRISPR/Cas9 knockout

To knock out TGFBR2, EGFR and AREG in PSCs, lenti-Cas9-Blast plasmids (52962; Addgene) were used. PSCs were infected and selected using 2 μg/mL blasticidin (A11139-03; Thermo Fisher Scientific). Single guide RNAs (sgRNA) were designed using Benchling and cloned into the LRGN (LentisgRNA-EFS-GFP-neo) plasmid. PSCs were plated as single clones in 96-well plates in the presence of geneticin (10131035; Thermo Fisher Scientific). Knockout was confirmed by western blot analysis or ELISA. sgRNAs against the *Rosa26* locus were included to generate control (i.e. WT) PSCs.

### RTK assay

PSCs were treated in Matrigel in 5%FBS DMEM with 20 ng/mL human TGFβ1 (T7039; Sigma) for 24 h. Phospho-RTK assays (ARY014; R&D Systems) were performed using 300 μg protein and following the Manufacturer’s instructions.

### Western blot analyses

PSCs and organoids were harvested in Cell Recovery Solution and incubated rotating for 30 minutes at 4C. Cells were pelleted and lysed in 0.1% Triton X-100, 15 mmol/L NaCl, 0.5 mmol/L EDTA, 5 mmol/L Tris, pH 7.5, supplemented with complete, mini protease inhibitors (11836170001; Roche) and a phosphatase inhibitor cocktail (4906837001; Roche). Cells were incubated on ice for 30 minutes before clarification. Standard procedures were used for western blotting. Primary antibodies used were HSP90α (07-2174; EMD Millipore), ACTIN (8456; Cell Signaling Technology), SMAD2 (5339; Cell Signaling Technology), pSMAD2/SMAD3 (8828; Cell Signaling Technology), TGFBR2 (AF532; R&D Systems), ERBB2 (2165; Cell Signaling Technology), p-ERBB2 (2243; Cell Signaling Technology), EGFR (4267; Cell Signaling Technology), p-EGFR (3777; Cell Signaling Technology), CC3 (9664; Cell Signaling Technology). Proteins were detected using HRP-conjugated secondary antibodies (Jackson ImmunoResearch Laboratories).

### ELISA assays

For ELISA of media, cultures were grown for 3 to 5 days. Media were collected and assayed using the manufacturer’s protocol. ELISA assays used were AREG (EMAREG; Thermo Fisher Scientific).

### Proliferation assays

For proliferation assays of PSCs in Matrigel, 5,000 PSCs were plated in 52 μL of 50% Matrigel in PBS on white 96-well plates (136101; Thermo Fisher Scientific) and cultured in 100 μL of media as specified in the figure legends. PSC proliferation was followed for 5 days with CellTiter-Glo (G7572; Promega) with measurements every 24 hours.

### Immunohistochemical and histological analyses

Standard procedures were used for IHC. Primary antibodies for IHC were p-EGFR (ab40815; Abcam), αSMA (ab5694; Abcam) and SV40 T antigen (ab16879, Abcam). Hematoxylin (H-3404, Vector Lab) was used as nuclear counterstain. Hematoxylin and eosin and Masson’s trichrome stains were performed according to standard protocols. Brightfield images of tissue slides were obtained with an Axio Vert.A1 (ZEISS). Quantification of metastatic areas over total areas in lung and liver tissues from NSG mice was done with QuPath software^42^. Stained sections were scanned with Aperio ScanScope CS and analyzed using the ImageScope Positive Pixel Count algorithm. The percentage of collagen area was then determined by calculating the percentage of blue pixels relative to the entire stained area. To quantify αSMA, p-EGFR and SV40 IHC, the percentage of strong positive pixels was calculated relative to the entire section with the ImageScope software.

### Flow cytometry

Tumors were processed as previously described^3^. For flow-cytometric analysis of myCAF/iCAF populations, cells were stained for 30 minutes with anti-mouse CD31-PE/Cy7 (102418; BioLegend), CD45-PerCP/Cy5.5 (103132; BioLegend), CD326 (EPCAM)-AlexaFluor 488, PDPN-APC/Cy7, MHCII-BV785 (107645, Biolegend), Ly6C-APC (128015; BioLegend) and CD90-PE (ab24904, Abcam) for 10 minutes with DAPI. For flow-cytometric analysis of immune cell populations, cells were stained for 30 minutes with anti-mouse CD45-PerCP/Cy5.5 (103132; BioLegend), TCR-β-Alexa488 (109215, Biolegend), CD3e-Alexa488 (100321, Biolegend), CD8-APC/Cy7 (100713, Biolegend), CD4-APC (100515, Biolegend), and for 10 minutes with DAPI.

### qPCR analyses

RNA (1 μg) was reverse transcribed using TaqMan reverse transcription reagents (N808-0234; Applied Biosystems). qPCR was performed using gene-specific TaqMan probes (Applied Biosystems) and master mix (4440040; Applied Biosystems). Gene expression was normalized to *Hprt*.

### RNA-sequencing and single-cell RNA-sequencing analyses

Samples were collected in 1 mL of TRIzol Reagent (15596018; Invitrogen). RNA was extracted using the PureLink RNA mini kit (12183018A; Invitrogen). RNA concentration was measured using a Qubit and RNA quality was assessed on a TapeStation 4200 (Agilent) using the Agilent RNA ScreenTape kit. mRNA library preparations were performed using 55 μL of 10 ng/mL per sample (RNA integrity number > 8). Illumina libraries were then sequenced on 1 lane of SP flowcell on NovaSeq6000. All RNA-seq data are available at the Gene Expression Omnibus (GEO) under the accession number GSE219180. Transcript counts were estimated using Salmon (version 1.4.0) against mouse reference genome GRCm38 (release 102) with default settings. Salmon estimated counts were summarized to gene level using the tximport package in RStudio for use with DESeq2. Protein coding genes with fewer counts than 2^5^ were filtered out before differential expression analysis (DEA). DEA was performed using DESeq package (V2) with default parameters in R. Genes with adjusted P < 0.05 were selected as significantly changed between conditions. GSEA was performed using the GSEA program (Broad Institute) on the Hallmark gene sets (h.all.v7.4) and the C2 canonical pathway collection (C2.all.v7.4) downloaded from the Molecular Signatures Database (MSigDB). Genes were ranked by their P values before submitted to GSEA for analysis. Heatmaps were plotted using Morpheus (Morpheus, https://software.broadinstitute.org/morpheus). The RNA-seq dataset of murine PDAC organoids is from Oni and Biffi et al^41^. Gene set variation analysis (GSVA)^43^ was performed on normalized gene expression using default parameters and the “gsva” method on available data from TCGA PAAD and TCGA BRCA. Correlation analyses were performed on z-scores of gene expression values or scaled GSVA scores of selected pathways using customized R scripts. The single-cell RNA-sequencing dataset of murine PDAC samples is from Elyada et al^6^.

### Statistical analysis

GraphPad Prism software, Morpheus software (Broad Institute), customized R scripts and Jupyter notebooks were used for graphical representation of data. Statistical analysis was performed using paired or unpaired Student’s t-test for or non-parametric Mann-Whitney test.

### Resource availability

Further information and requests for resources and reagents should be directed to and will be fulfilled by the lead contact, Giulia Biffi.

All unique/stable reagents generated in this study are available from the lead contact with a completed Materials Transfer Agreement.

## SUPPLEMENTARY FIGURE LEGENDS

**Figure S1. TGF-β and PDAC organoid-conditioned media induce ERBB activation in myCAFs. Related to Figures 1 and 2. (A)** Western blot analysis of p-EGFR, EGFR, p-ERBB2 and ERBB2 in human pancreatic stellate cells (PSCs) cultured for 24 h in Matrigel in control media (i.e. 5% FBS DMEM) with or without 20 ng/mL TGF-β. ACTIN, loading control. **(B)** Heatmap of scaled expression of *Egfr* and *Erbb2* in different cell populations of pancreatic tumors of the KPC (*Kras*^LSL-G12D/+^; *Trp53*^LSL-R172H/+^; *Pdx1-Cre*) mouse model of PDAC (n=4), as analyzed by single-cell RNA-sequencing (scRNA-seq). Data are scaled such that the cluster with the lowest average expression = 0 and the highest = 1 for each gene. The dataset analyzed is from Elyada et al.^6^. **(C)** Heatmap of scaled expression of *EGFR* and *ERBB2* in different cell populations of human PDAC tumors (n=6), as analyzed by scRNA-seq. Data are scaled such that the cluster with the lowest average expression = 0 and the highest = 1 for each gene. The dataset analyzed is from Elyada et al.^6^. **(D-F)** Validation of *Tgfbr2* KO PSCs. **(D)** Western blot analysis of TGFBR2 in murine *Tgfbr2* wild-type (WT) and knock out (KO) (2 PSC lines, 5 clones from 3 different guide RNAs) PSCs cultured in monolayer in control media. ACTIN, loading control. **(E)** qPCR analysis of TGF-β signaling targets (*Ctgf, Col1a1, Tgfb1*) in murine *Tgfbr2* WT and KO PSCs cultured for 4 days in Matrigel in control media with or without 20 ng/mL TGF-β. Results show mean ± standard error of mean (SEM) of 2-5 biological replicates per group. *, *P* < 0.05; ***, *P* < 0.001, paired Student’s t-test. **(F)** Proliferation curves of murine *Tgfbr2* WT and KO PSCs cultured for 5 days in Matrigel in control media with or without 20 ng/mL TGF-β. Results show mean ± standard deviation (SD) of 5 technical replicates per cell line. ***, P < 0.001, unpaired Student’s t-test calculated for the last time point. **(G)** Western blot analysis of p-EGFR, EGFR, p-SMAD2 and SMAD2 in murine PSCs cultured for 4 days in Matrigel in control media with or without 20 ng/mL TGF-β in the presence or absence of 2 μM A83-01 (TGFBR1 inhibitor, TGFBRi). ACTIN, loading control. **(H)** Heatmap of scaled expression of *Tgfb1* in different cell populations of pancreatic tumors of the KPC mouse model of PDAC (n=4), as analyzed by scRNA-seq. Data are scaled such that the cluster with the lowest average expression = 0 and the highest = 1 for each gene. The dataset analyzed is from Elyada et al.^6^. **(I)** Heatmap of scaled expression of *TGFB1* in different cell populations of human PDAC tumors (n=6), as analyzed by scRNA-seq. Data are scaled such that the cluster with the lowest average expression = 0 and the highest = 1 for each gene. The dataset analyzed is from Elyada et al.^6^. **(J)** RNA-seq expression of *Tgfb1* in murine PDAC organoids derived from the KPC mouse model (n=21). Data are from Oni and Biffi et al.^41^. **(K)** GSEA of ERBB signature (i.e. 138 genes from 2B) in PSCs cultured for 4 days in Matrigel in control media with 20 ng/mL TGF-β compared to control PSCs. **(L)** GSEA of cholesterol biosynthesis in PSCs cultured for 4 days in Matrigel in control media with 20 ng/mL TGF-β compared to control PSCs. **(M)** Heatmap showing GSVA scores of pathways significantly positively correlated with the human myCAF signature in TCGA PAAD (n=168). The human myCAF signature is from Elyada et al.^6^. GSVA scores were scaled as z-scores.

**Figure S2. A TGF-β-induced autocrine amphiregulin signaling mediates EGFR activation in myCAFs. Related to Figure 3. (A)** qPCR analysis of ERBB ligands (*Hbegf, Areg, Ereg, Btc, Egf, Tgfa, Nrg1*) and TGF-β signaling targets (*Col1a1, Ctgf, Tgfb1*) in PSCs cultured for 10 min, 30 min, 1 h or 24 h in Matrigel in control media with 20 ng/mL TGF-β. Results show mean ± SEM of 4-6 biological replicates. *, *P* < 0.05; **, *P* < 0.01, ***, *P* < 0.001, paired Student’s t-test. **(B)** qPCR analysis of *Hbegf* in murine control (i.e unmodified), WT (i.e. *Rosa26* KO), *Tgfbr2* KO or *Egfr* KO PSCs cultured for 4 days in Matrigel in control media with or without 20 ng/mL TGF-β in the presence or absence of 1 μM Erlotinib (EGFRi), 300 nM Neratinib (ERBBi) or 2 μM A83-01 (TGFBRi). Results show mean ± SEM of 3-11 biological replicates. *, *P* < 0.05; **, *P* < 0.01, ***, *P* < 0.001, paired and unpaired Student’s t-test. **(C-F)** Validation of *Egfr* KO PSCs. **(C)** Western blot analysis of EGFR in murine *Egfr* WT and KO PSCs (2 PSC lines, 5 clones from 3 different guide RNAs) cultured in monolayer in control media. ACTIN, loading control. **(D)** Bright field images of murine *Egfr* WT and KO PSCs cultured for 5 days in Matrigel in control media with or without 10 ng/mL EGF. Scale bars, 100 μm. **(E)** Proliferation curves of murine *Egfr* WT and KO PSC4 cultured for 5 days (120 h) in Matrigel in control media with 20 ng/mL TGF-β. Results show mean ± SD of 5 technical replicates per cell line. **, *P* < 0.01, ***, *P* < 0.001, unpaired Student’s t-test calculated for the last time point. **(F)** Proliferation curves of murine *Egfr* WT and KO PSC5 cultured for 5 days (120 h) in Matrigel in control media with 20 ng/mL TGF-β. Results show mean ± SD of 5 technical replicates per cell line. ***, *P* < 0.001, unpaired Student’s t-test calculated for the last time point. **(G)** Western blot analysis of p-EGFR, EGFR, p-ERBB2, ERBB2, p-SMAD2 and SMAD2 in murine PSCs cultured for 4 days in control media with or without 20 ng/mL TGF-β in the presence or absence of 1 μM Erlotinib (EGFRi) or 300 nM Neratinib (ERBBi). ACTIN, loading control. **(H)** Spearman’s correlation coefficients (R) between normalized gene expression of *TGFB1* and normalized gene expression of myCAF markers (*CTGF*, *GLI1, COL1A1* and *ACTA2*), *HBEGF* and *AREG* from human PDAC samples analyzed by bulk RNA-seq. Correlation analyses were performed on z-scores of gene expression values. Statistically significant correlations were only considered when *P* < 0.05. Data are from TCGA PAAD (n=168). **(I)** ELISA of AREG from media of murine *Areg* WT and KO PSCs cultured for 4 days in Matrigel in control media with or without 20 ng/mL TGF-β. Results show mean ± SEM of 2-3 biological replicates, respectively. **, *P* < 0.01; ***, *P* < 0.001, paired Student’s t-test. **(J)** Western blot analysis of p-EGFR and EGFR in murine *Areg* WT and KO PSCs cultured for 24 h in Matrigel in control media with or without 20 ng/mL TGF-β. ACTIN, loading control.

**Figure S3. Inhibition of ERBB signaling impairs myCAF proliferation and signature *in vitro*. Related to Figure 4. (A)** Proliferation curves of murine PSCs cultured for 5 days (120 h) in Matrigel in control media with or without 20 ng/mL TGF-β in the presence or absence of 1 μM Erlotinib (EGFRi). Results show mean ± SD of 5 technical replicates. *, *P* < 0.05, ***, *P* < 0.001, unpaired Student’s t-test calculated for the last time point. **(B)** Western blot analysis of p-EGFR, EGFR, p-ERBB2 and ERBB2 in PSCs cultured for 4 days in Matrigel in control media, PDAC organoid CM or CM in the presence of 300 nM Neratinib (ERBBi) from day 0 (d0) or from day 3 for the last 24 h (d3). HSP90 and ACTIN, loading controls. **(C)** Proliferation curves of murine PSCs cultured for 5 days (120 h) in Matrigel in control media, in PDAC organoid CM or in CM in the presence of 300 nM Neratinib (ERBBi) starting from 72 h for the last 48 h. Results show mean ± SD of 5 technical replicates. ***, *P* < 0.001, unpaired Student’s t-test calculated for the last time point. **(D)** Proliferation curves of murine PSCs cultured for 5 days (120 h) in Matrigel in control media or in PDAC organoid CM in the presence or absence of 1 μM Erlotinib (EGFRi). Results show mean ± SD of 5 technical replicates. **, *P* < 0.01; ***, *P* < 0.001, unpaired Student’s t-test calculated for the last time point. **(E)** GSEA of apoptosis signaling in PSCs cultured for 4 days in Matrigel in PDAC organoid CM with ERBBi compared to PSCs cultured in CM. The signature was not significantly altered. **(F)** Western blot analysis of p-EGFR and cleaved caspase 3 (CC3) in PSCs cultured for 4 days in Matrigel in control media, PDAC organoid CM or CM in the presence of 300 nM Neratinib (ERBBi). HSP90, loading control. **(G)** Selected pathways found significantly enriched or depleted (FDR < 0.25) by GSEA in PSCs cultured for 4 days with PDAC organoid CM following treatment with the JAK inhibitor (JAKi) AZD1480, which targets iCAFs^3^. The RNA-seq dataset analyzed is from Biffi et al.^3^. **(H)** Pathways found significantly enriched (FDR < 0.05) by DAVID analysis following ERBB inhibition in PSCs cultured for 4 days with PDAC organoid CM, as assessed by RNA-seq. **(I)** Bright field images of KPC PDAC organoids cultured for 4 days in control media with or without 300 nM ERBBi. Scale bars, 500 μm.

**Figure S4. Inhibition of ERBB signaling targets myCAFs *in vivo*. Related to Figure 5. (A)** Representative hematoxylin and eosin (H&E) stain in 2-week vehicle- and ERBBi-treated orthotopically grafted organoid-derived mouse models. Scale bars, 200 μm. **(B)** Representative p-EGFR IHC in 2-week vehicle- and ERBBi-treated tumors. Scale bars, 50 μm. **(C)** Quantification of p-EGFR stain in 2-week vehicle- (n=11) and ERBBi- (n=9) treated tumors. Results show mean ± SEM. *, *P* < 0.05, Mann-Whitney test. **(D)** Flow cytometric analysis of immune cells (CD45^+^ CD31^-^) from live singlets in vehicle- (n=11) and ERBBi- (n=8) treated tumors. Results show mean ± SEM. No statistical difference was found, as calculated by Mann-Whitney test. **(E)** Flow cytometric analysis of total T cells, CD4^+^ T cells and CD8^+^ T cells from live singlets in vehicle- (n=11) and ERBBi- (n=8) treated tumors. Results show mean ± SEM. *, *P* < 0.05; **, *P* < 0.01, Mann-Whitney test. **(F)** Flow cytometric analysis of endothelial cells (CD31^+^CD45^-^), epithelial cells (CD45-CD31-EpCAM^+^) and CAFs (CD45^-^CD31^-^EpCAM^-^PDPN^+^) from live singlets in vehicle- (n=11) and ERBBi- (n=8) treated tumors. Results show mean ± SEM. No statistical difference was found, as calculated by Mann-Whitney test. **(G)** Schematic of flow cytometric strategy of PDAC CAF subtypes from 2-week vehicle- and ERBBi-treated orthotopically grafted PDAC tumors. **(H)** UMAP plots of the fibroblast cluster from PDAC tumors from KPC mice showing the iCAF, myCAF and apCAF clusters (left) and *Thy1* (CD90) expression (right). scRNA-seq datasets were obtained from Elyada et al.^6^. **(I)** myCAF/(iCAF+apCAF) ratio from live singlets in vehicle- (n=11) and ERBBi- (n=8) treated tumors. Results show mean ± SEM. *, *P* < 0.05, Mann-Whitney test.

**Figure S5. ERBB-activated myCAFs promote local metastasis of PDAC. Related to Figure 6. (A)** Representative H&E stain in tumors derived from the transplantation of PDAC organoids with or without *Egfr* WT or *Egfr* KO PSCs. Scale bars, 200 μm. **(B)** Representative SV40 IHC in tumors derived from the transplantation of PDAC organoids with or without *Egfr* WT or *Egfr* KO PSCs. Scale bars, 50 μm. **(C)** Quantification of SV40 T antigen stain in tumors derived from the transplantation of PDAC organoids with or without *Egfr* WT or *Egfr* KO PSCs. Results show mean ± SEM of 23-25 biological replicates. ***, *P* < 0.001, Mann-Whitney test. **(D)** Representative Masson’s trichrome stain in tumors derived from the transplantation of PDAC organoids with or without *Egfr* WT or *Egfr* KO PSCs. Scale bars, 50 μm. **(E)** Quantification of Masson’s trichrome stain in tumors derived from the transplantation of PDAC organoids with or without *Egfr* WT or *Egfr* KO PSCs. Results show mean ± SEM of 23-25 biological replicates. No statistical difference was found, as calculated by unpaired Student’s t-test. **(F)** Representative αSMA IHC in tumors derived from the transplantation of PDAC organoids with or without *Egfr* WT or *Egfr* KO PSCs. Scale bars, 50 μm. **(G)** Quantification of αSMA stain in tumors derived from the transplantation of PDAC organoids with or without *Egfr* WT or *Egfr* KO PSCs. Results show mean ± SEM of 23-25 biological replicates. No statistical difference was found, as calculated by unpaired Student’s t-test. **(H)** Flow cytometric analyses of immune cells (CD45^+^CD31^-^), endothelial cells (CD31^+^CD45^-^), epithelial cells (CD31-CD45-EpCAM^+^) and CAFs (CD45^-^CD31^-^EpCAM^-^PDPN^+^) from live singlets in tumors derived from the transplantation of PDAC organoids with or without *Egfr* WT or *Egfr* KO PSCs. Results show mean ± SEM (n=19-20/cohort). *, *P* < 0.05, unpaired Student’s t-test. **(I)** myCAF/iCAF ratio from live singlets in tumors derived from the transplantation of PDAC organoids with or without *Egfr* WT or *Egfr* KO PSCs. Results show mean ± SEM (n=19-20/cohort). *, *P* < 0.05, Mann-Whitney test. **(J)** Quantification of metastatic burden (metastatic area/total area) in liver tissues of mice transplanted with PDAC organoids with or without *Egfr* WT or *Egfr* KO PSCs (n=24-25 mice/cohort). No statistical difference was found, as calculated by Mann-Whitney test.

**Figure S6. EGFR activation occurs in myofibroblastic CAFs in various malignancies. (A)** Heatmap showing GSVA scores of pathways positively correlated with the myofibroblastic matrix CAF (mCAF) signature in TCGA breast cancer BRCA (n=1100). The mCAF signature was derived from the murine breast cancer scRNA-seq dataset of Bartoschek et al.^18^ and converted into human genes prior analysis. GSVA scores were scaled as z-scores. **(B)** Spearman’s correlation coefficients (R) between normalized gene expression of *TGFB1* and normalized gene expression of *COL1A1, ACTA2, CTGF, GLI1* and *AREG* from tumor samples analyzed by bulk RNA-seq. Correlation analyses were performed on z-scores of gene expression values. Statistically significant correlations were only considered when *P* < 0.05. Data are from TCGA breast cancer BRCA (n=1100). **(C)** Spearman’s correlation coefficients (R) between normalized gene expression of *TGFB1* and normalized gene expression of *COL1A1, ACTA2, CTGF, GLI1* and *AREG* from tumor samples analyzed by bulk RNA-seq. Correlation analyses were performed on z-scores of gene expression values. Statistically significant correlations were only considered when *P* < 0.05. Data are from TCGA lung cancer LUAD (n=518). **(D)** qPCR analysis of *Areg* in murine pulmonary fibroblasts cultured for 4 days in Matrigel in control media with or without 20 ng/mL TGF-β. Results show mean ± SD of 5 technical replicates. *, *P* < 0.05, paired Student’s t-test. **(E)** Western blot analysis of p-EGFR and EGFR in mouse pulmonary fibroblasts (MPFs) cultured for 4 days in Matrigel in control media with or without 20 ng/mL TGF-β. ACTIN, loading control. **(F)** qPCR analysis of *Dusp6* in murine pulmonary fibroblasts cultured for 4 days in Matrigel in control media with or without 20 ng/mL TGF-β. Results show mean ± SD of 5 technical replicates. *, *P* < 0.05, paired Student’s t-test.

