## Supplementary figures and images for "ERBB-activated myofibroblastic cancer-associated fibroblasts promote local metastasis of pancreatic cancer"

### Supplemental Figure S1

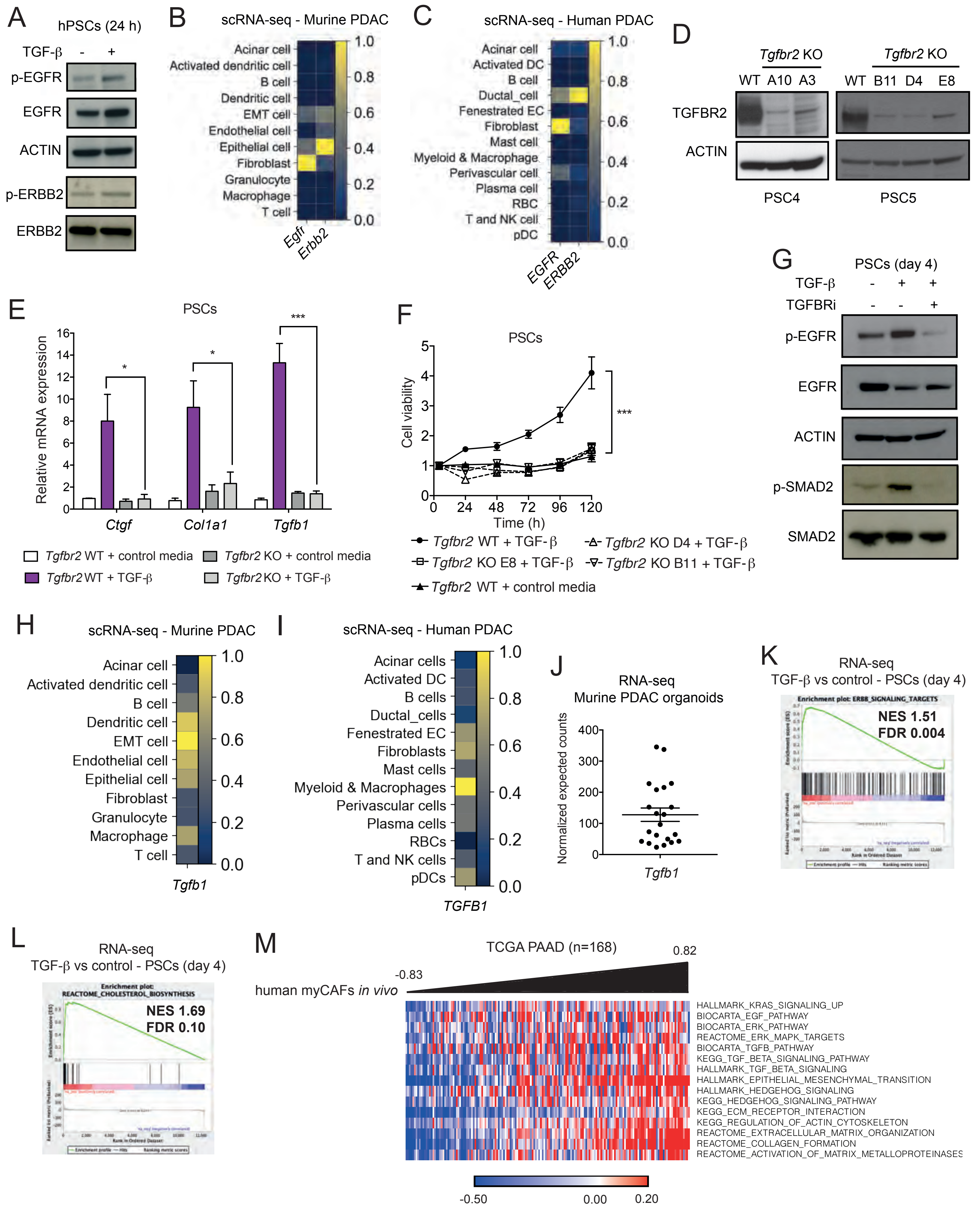

### Supplemental Figure S2

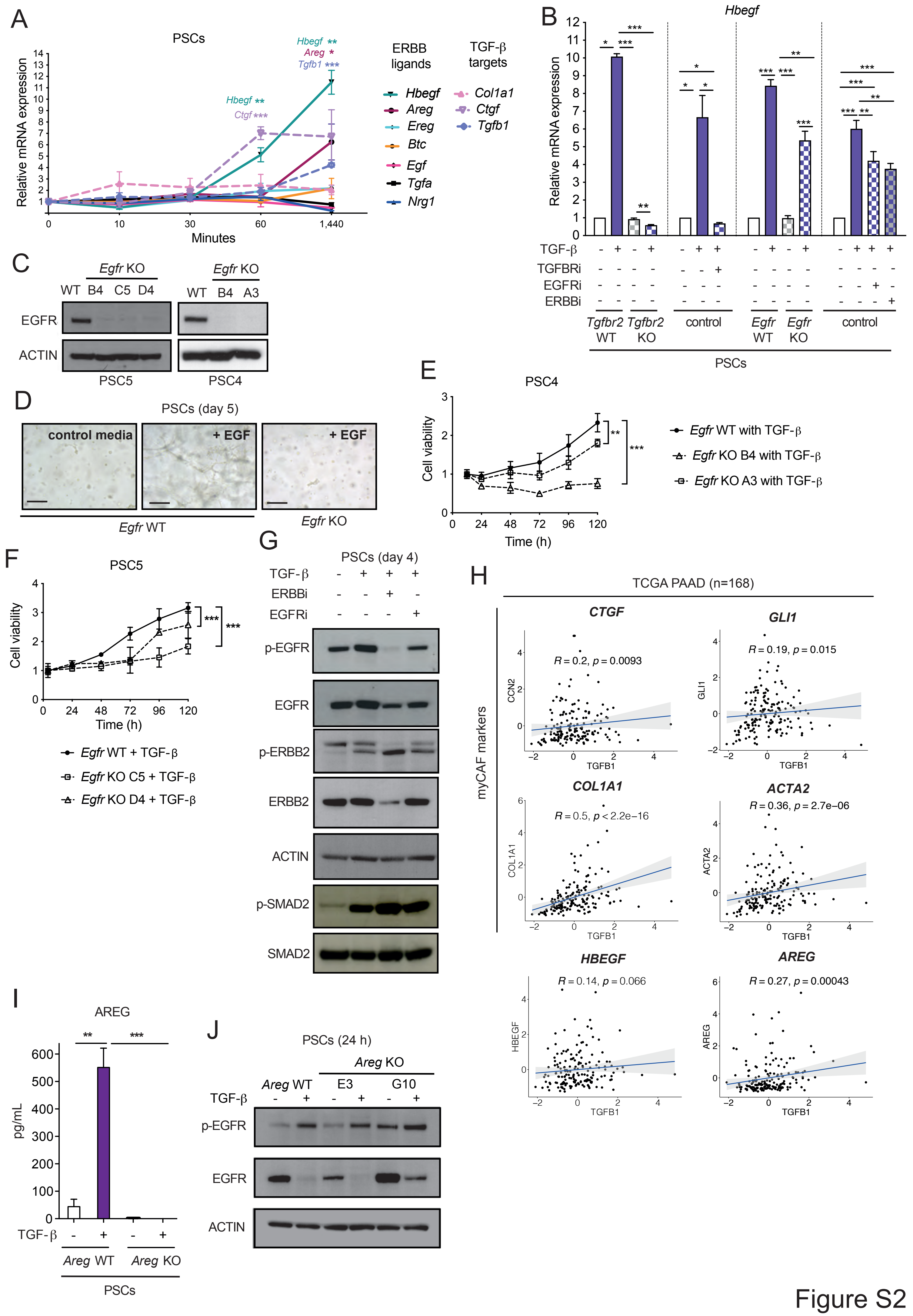

### Supplemental Figure S3

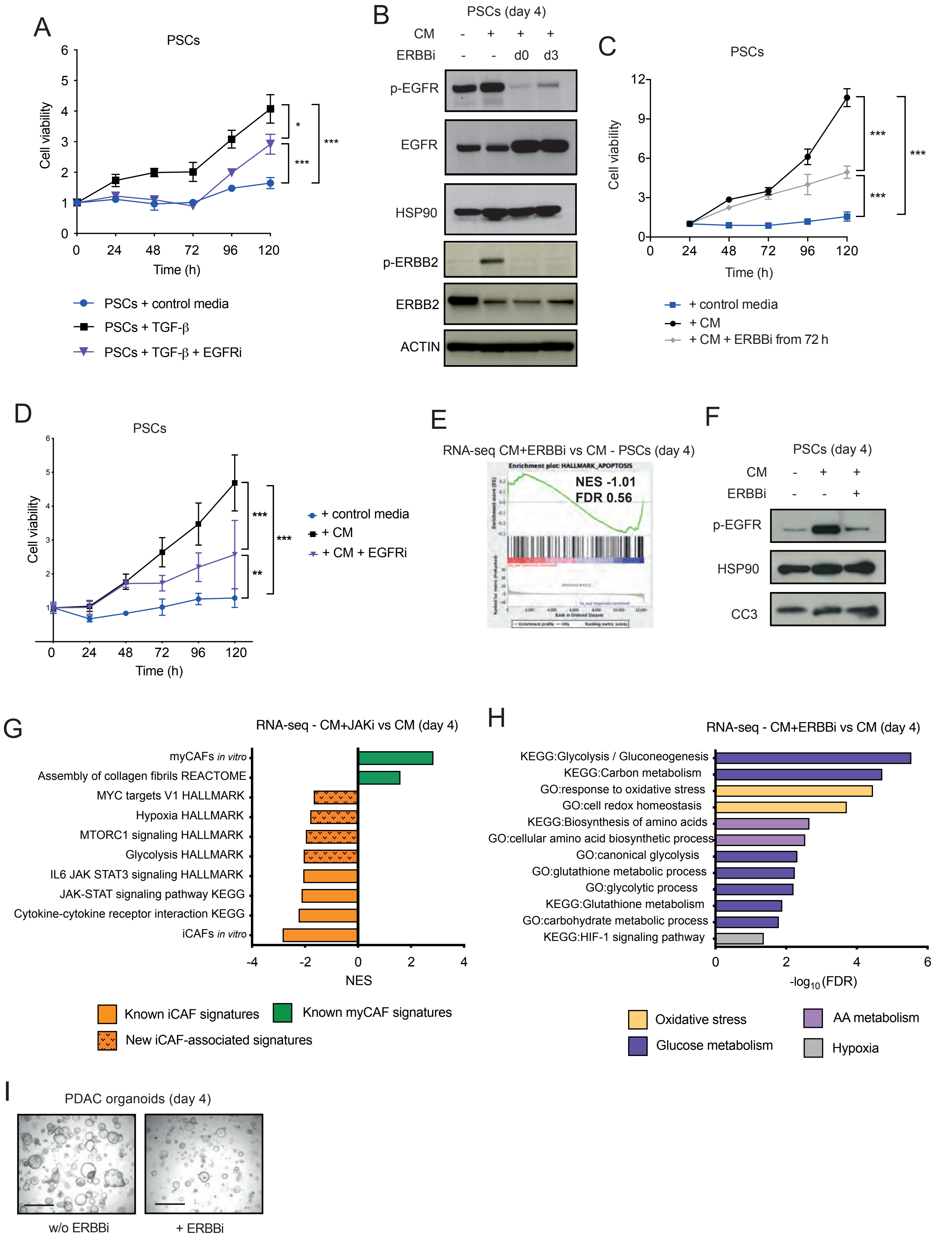

### Supplemental Figure S4

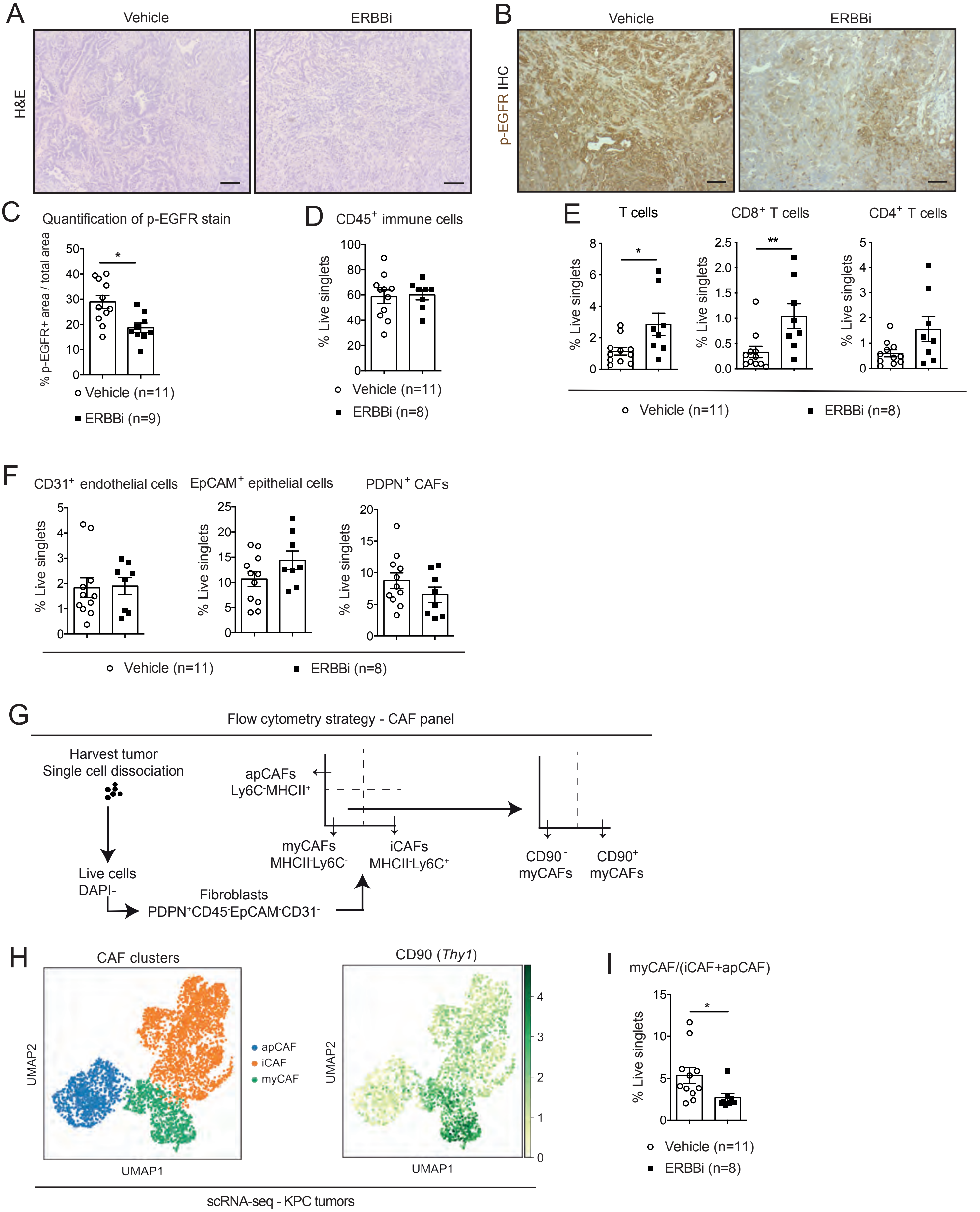

### Supplemental Figure S5

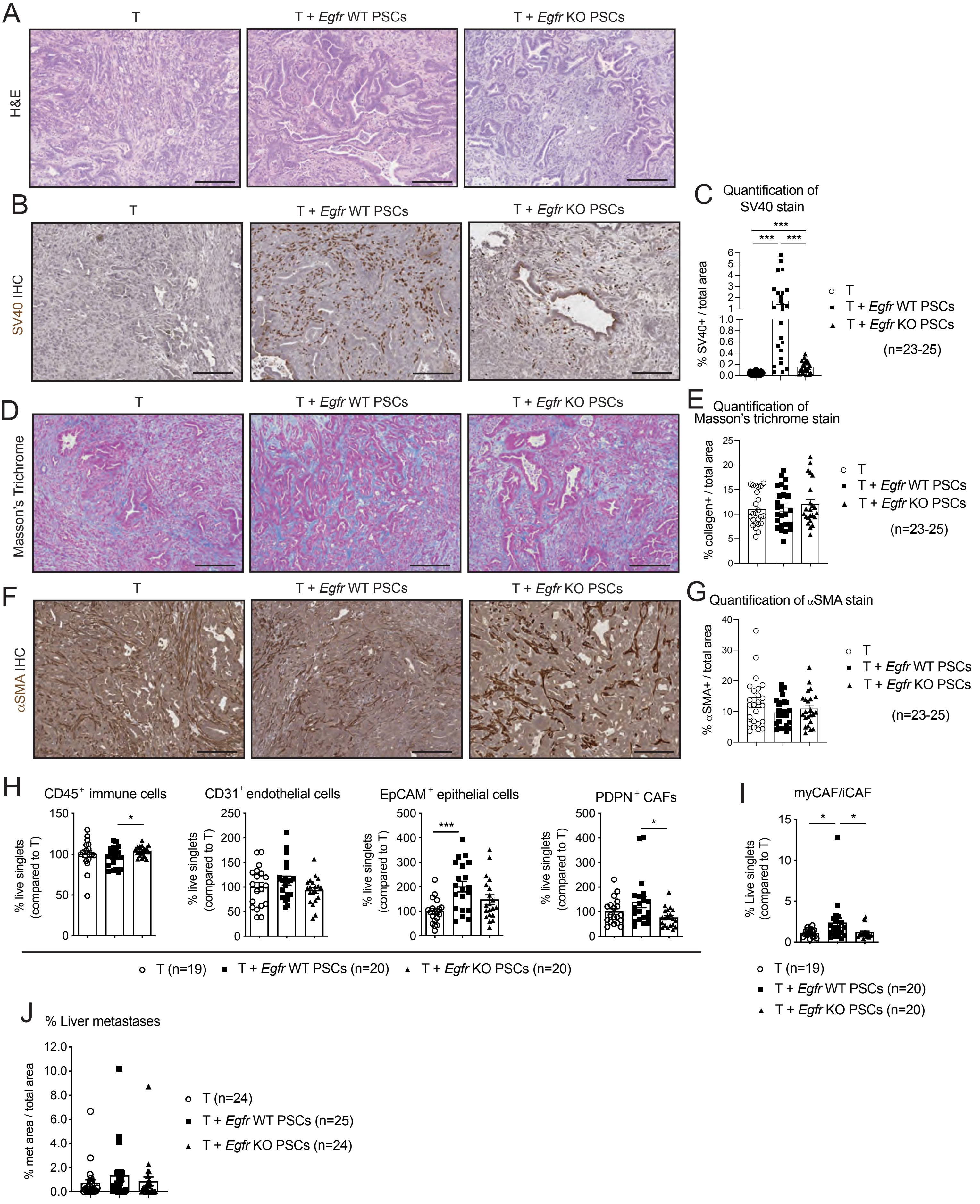

### Supplemental Figure S6

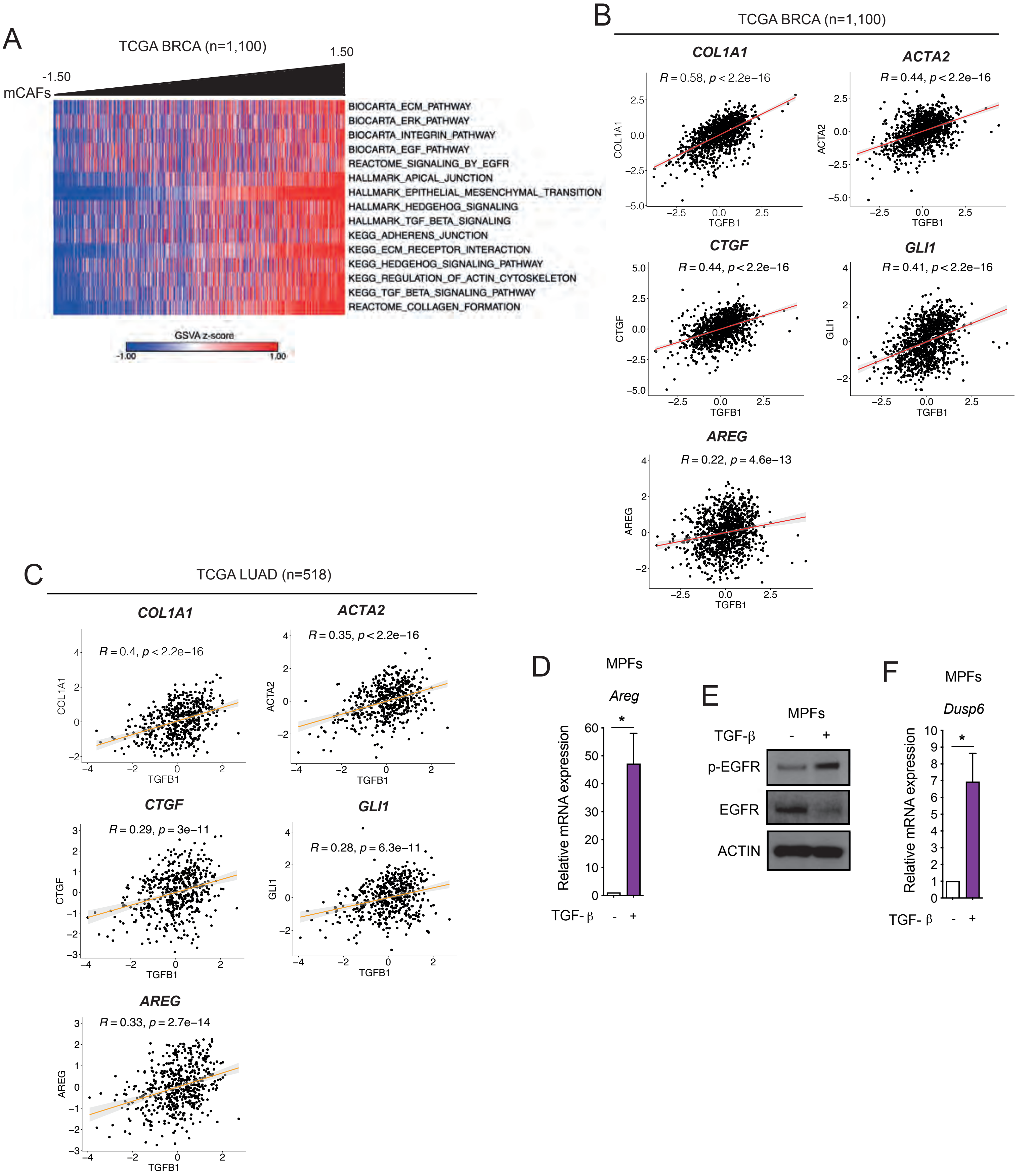
